# *In vitro* Characterization of Peptidomimetic Proteolysis Targeting Chimera (PROTAC) as a Degrader of 3-Chymotrypsin-Like Protease (Mpro/3CLpro) against SARS-CoV-2

**DOI:** 10.1101/2025.10.30.685646

**Authors:** Mirko G. Liturri, A. Bergna, A. Lai, C. Della Ventura, A. Gabrieli, I. Seravalli, S. Ciofi-Baffoni, E. Lenci, A. Trabocchi, S. Rusconi

**Author notes:** **Corresponding Author** Stefano Rusconi.

## Abstract

The SARS-CoV-2 main protease (3CLpro) is a key target for antiviral development. We investigated FT235, a peptidomimetic PROTAC linking a GC-376 warhead to pomalidomide for targeted degradation. FT235 bound 3CLpro, inhibiting activity (IC_50_ = 21.2 µM), and reducing protease levels in cells. *In vitro* data showed no cytotoxicity up to 100 µM and variant-dependent antiviral activity, with increased potency in the presence of a P-gp inhibitor. These results support PROTAC-based antivirals as promising therapeutic candidates.

## Main Text

The SARS-CoV-2 main protease (Mpro/3CLpro) is essential for viral maturation and replication, representing a major target for antiviral development. Clinically approved drugs such as nirmatrelvir/ritonavir are effective but limited by drug–drug interactions (1) and frequent dosing requirements (2). To explore alternative strategies, a chemoinformatic screening of 250 compounds identified FT235, a peptidomimetic PROTAC designed to degrade 3CLpro. FT235 links a GC-376-derived dipeptidyl ligand to a pomalidomide moiety via a piperazine–piperidine linker, enabling targeted proteasomal degradation (3).

X-ray crystallography and NMR confirmed that FT235 binds the catalytic site of 3CLpro, validating engagement of the dipeptidyl warhead with the active cysteine and supporting potential ubiquitin ligase recruitment. FRET-based enzymatic assays using the Hilyte Fluor-488-ESATLQSGLRKAK-(QXL-520)-NH_2_ substrate (60 nM) demonstrated measurable inhibitory activity, with an IC_5 0_ of 21.2 ± 5.8 μM. Although weaker than covalent inhibitors, FT235 functions as both an enzymatic inhibitor and a degrader, as shown in cell-based assays. Intracellular 3CLpro levels decreased significantly, consistent with GC-376-based binding recruiting cereblon E3 ligase through the pomalidomide unit, leading to ubiquitination and proteasomal clearance (3).

Based on these preliminary data, we next investigated the *in vitro* cytotoxicity profile of FT235. For this purpose, VERO E6 cells [also known as Vero C1008, obtained from the American Type Culture Collection, ATCC CRL-1586, Manassas, VA, USA] were employed. Cells were seeded at a density of 1 × 10^5 cells/well and pre-incubated for 24 h before treatment. Serial two-fold dilutions of FT235 were prepared, starting from a maximum concentration of 200 μM. Cell viability was assessed using the luminescent CellTiter-Glo 2.0 assay (Promega, Madison, WI, USA) following the manufacturer’s instructions, and luminescence was recorded using a GloMax® Discover multimode microplate reader [REF GM3000 (Promega Corporation, USA)].

The 50% cytotoxic concentration (CC_5 0_) was determined by nonlinear dose– response curve fitting using GraphPad Prism (v.9.0.2). Untreated cells and DMSO-treated cells were used as 0% and 100% viability controls, respectively, to normalize the data. Each compound concentration was tested in at least three technical replicates across a minimum of three independent experiments. Notably, cytotoxicity assays were performed in the presence and absence of the P-glycoprotein (P-gp) inhibitor CP-100356 monohydrochloride (CAS n° 142715-48-8) at 0.5 µM, a concentration corresponding to its reported IC_5 0_ for MDR1 inhibition (4). The results showed no detectable cytotoxicity at concentrations up to 100 µM, supporting a low cytotoxic profile for FT235 under the tested conditions (Fig. 1).

**Fig 1.**
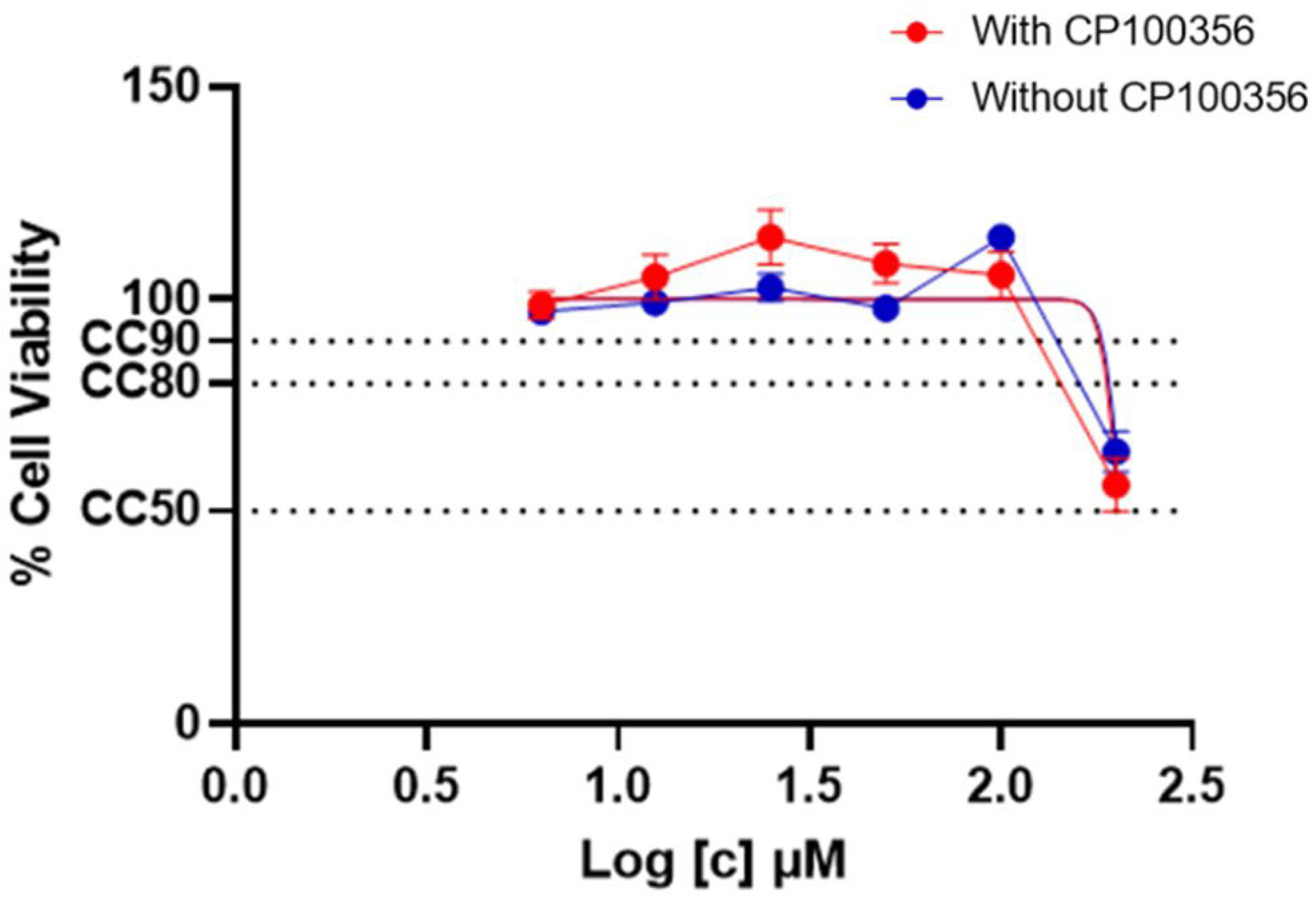
FT235 cytotoxocity with and without P-gp inhibitor.

The antiviral activity of FT235 was assessed in VERO E6 cells infected with SARS-CoV-2 variants B.1 [EPI_ISL_2472896], BA.5 [EPI_ISL_18450047], and JN.1 [EPI_ISL_18863880].

Swabs positivity were evaluated with the multiplexed RT-qPCR developed by English consortium (https://www.protocols.io/view/multiplexed-rt-qpcr-to-screen-for-sars-cov-2-b-1-1-br9vm966?). Whole genome sequences were obtained using a modified version of the ARTIC protocol (https://artic.network/ncov-2019; accessed September 29, 2025) using the Illumina DNA Prep and the IDT ILMN DNA/RNA Index kit (Illumina, San Diego, CA, USA). Sequencing was performed on Illumina Miseq platform (Illumina Reads were mapped and aligned to the reference genome obtained from GISAID (https://www.gisaid.org/) [NC_045512.2] using Geneious Prime software v. 2025.2 (Biomatters, Auckland, New Zealand) (http://www.geneious.com). Obtained sequences were classified using the Pangolin COVID-19 Lineage Assigner v. 4.3 (https://pangolin.cog-uk.io/) and Nextclade v. 2.14.1 (https://clades.nextstrain.org/).

Cells were pre-incubated for 1 h with two-fold serial dilutions of FT235, followed by infection at MOI 0.01 and 72 h incubation. Assays were performed with or without the P-glycoprotein (P-gp) inhibitor CP100356 (0.5–1.5 µM) to assess the contribution of efflux mechanisms. FT235 showed moderate activity against the ancestral B.1 strain (IC_5 0_ = 187.99 ± 20.78 µM), which improved to 77.52 ± 3.21 µM in the presence of CP100356. Significant differences were observed at 100, 50, and 25 µM (p < 0.0001–0.0068), whereas no differences were detected at lower concentrations.

FT235 efficacy was partially reduced at 1.0–1.5 µM CP100356 compared to 0.5 µM, although most differences remained statistically significant (p < 0.0001–0.0156). (IC_5 0_ = 96.13–120.06 µM), although differences remained significant in most cases (p < 0.0001–0.0156) (Fig. 2A). Based on these results, 0.5 µM CP100356 was selected for subsequent experiments, as this concentration showed the highest cell viability at the first FT235 concentration (100 µM) compared to 1.0 and 1.5 µM CP100356 (statistically significant for both), while no significant differences were observed at lower FT235 concentrations. This allowed the selection of a P-gp inhibitor concentration that optimized cell viability across the full range of FT235 concentrations (Fig. 2B).

**Fig 2.**
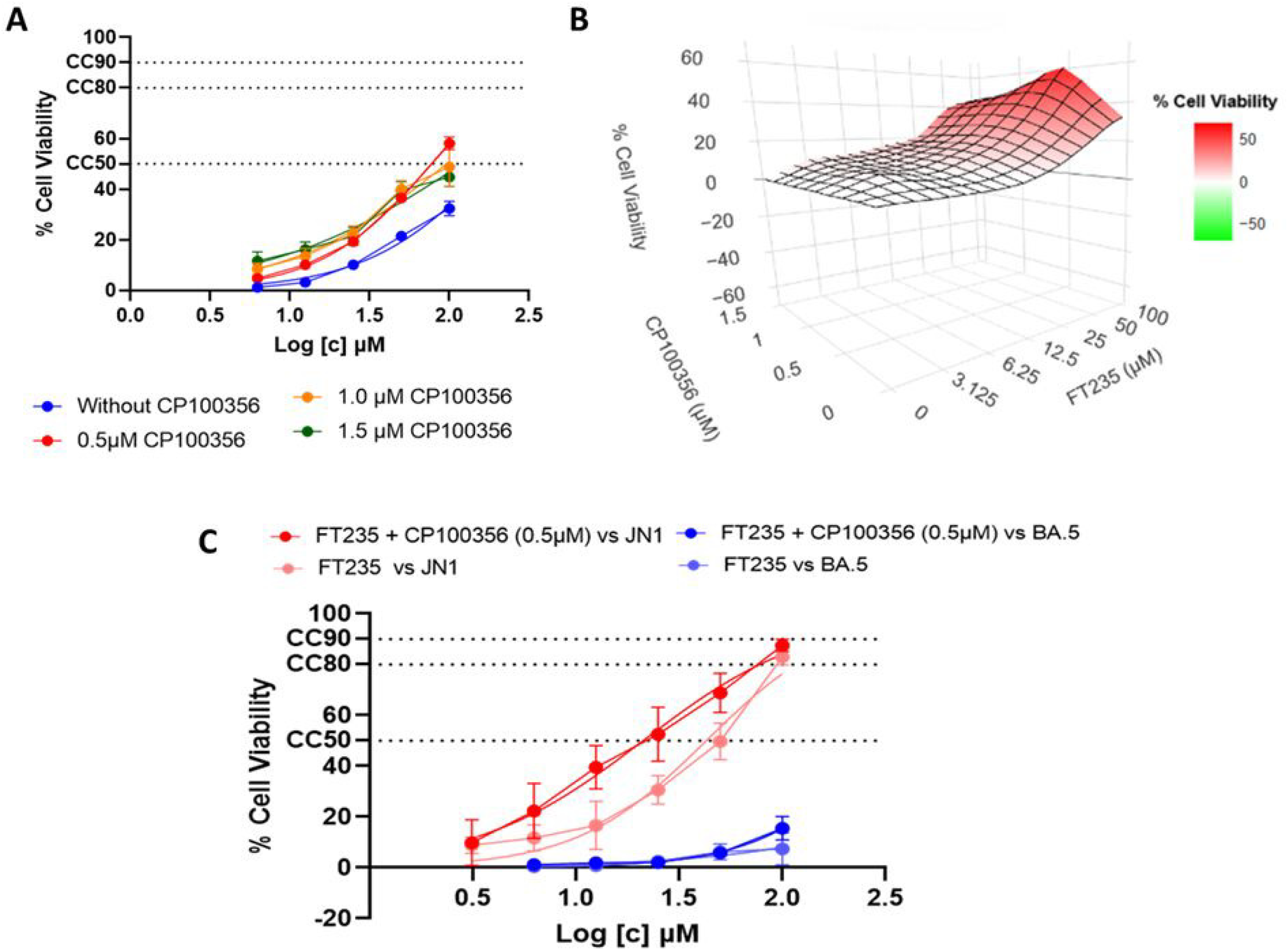
Antiviral efficacy of FT235. (A) Antiviral activity of FT235 against SARS-CoV-2 B.1 variant. (B) 3D representation of the interaction between FT235 and CP100356 on cell viability, highlighting the optimal dosage of the P-gp inhibitor to maximize the response. (C) Comparison of FT235 antiviral activity against JN.1 and BA.5 with or without P-gp inhibitor

Against JN.1, FT235 showed greater potency (IC_5 0_ = 44.27 ± 4.99 µM without and 18.08 ± 4.08 µM with CP100356), consistent with an enhanced effect upon P-gp inhibition. In contrast, no measurable activity was detected against the Omicron BA.5 variant, suggesting that structural changes in 3CLpro may impair PROTAC efficacy (Fig. 2C). Focusing on the nsp5 region encoding 3CLpro, catalytic residues were highly conserved across B.1, BA.5, and JN.1, with no significant amino acid changes. Nevertheless, mutations in distal or allosteric regions could subtly affect protease conformation and active site accessibility. These structural variations likely contribute to the observed differences in FT235 antiviral activity, with higher potency against JN.1 and reduced efficacy against BA.5 indicating that PROTAC-mediated degradation depends not only on catalytic site conservation but also on the overall 3CLpro structure and warhead engagement.

These results indicate that FT235 represents a promising candidate for the selective degradation of 3CLpro, exhibiting low cytotoxicity and variant-dependent antiviral activity. The limited efficacy observed against BA.5 highlights the need for structural optimization to accommodate emerging mutations in 3CLpro.

Overall, our findings support peptidomimetic PROTACs as a versatile platform for next-generation antiviral development. These preliminary observations underscore how PROTAC technology, already widely employed in oncology for the development of targeted antineoplastic therapies against oncoproteins (5), could also be applied in the context of infectious diseases, opening new avenues for the development of innovative molecules. These results further motivate investigations into pharmacokinetics, in vivo efficacy, and broad-spectrum activity.

## Acknowledgments

Supported by Intesa Sanpaolo (B/2021/0212, B/2022/0209) EU NextGenerationEU– PNRR (THE, ECS00000017, CUP B83C22003920001), MUR Departments of Excellence (DICUS 2.0), UniFI Chemistry “Ugo Schiff” and an unrestricted research grant from Fondazione Sodalitas, Milan, to SR.

## References

1. Girardin F, Manuel O, Marzolini C, Buclin T. 2022. Evaluating the risk of drug-drug interactions with pharmacokinetic boosters: the case of ritonavir-enhanced nirmatrelvir to prevent severe COVID-19. Clin Microbiol Infect 28:1044–1046.

2. Lange NW, Salerno DM, Jennings DL, Choe J, Hedvat J, Kovac DB, Scheffert J, Shertel T, Ratner LE, Brown RS Jr, Pereira MR. 2022. Nirmatrelvir/ritonavir use: Managing clinically significant drug-drug interactions with transplant immunosuppressants Am J Transplant 22:1925–1926.

3. Grifagni D, Lenci E, De Santis A, Orsetti A, Barracchia CG, Tedesco F, Bellini Puglielli R, Lucarelli F, Lauriola A, Assfalg M, Cantini F, Calderone V, Guardavaccaro D, Trabocchi A, D’Onofrio M, Ciofi-Baffoni S. 2024. Development of a GC-376 Based Peptidomimetic PROTAC as a Degrader of 3-Chymotrypsin-like Protease of SARS-CoV-2. ACS Med Chem Lett 15:250–257.

4. https://www.medchemexpress.com/cp-100356-hydrochloride.html?srsltid=AfmBOopbB7qm94dvOO_Z7PItvM8rESWiOhxjKiMl_RQgjTyZMQZdBpJd

5. Li X, Song Y. 2020. Proteolysis-targeting chimera (PROTAC) for targeted protein degradation and cancer therapy. J Hematol Oncol 13:50.

